# Selectivity Filter Dynamics Define Ion Conductance and Selectivity Differences in CNG and HCN Channels

**DOI:** 10.1101/2025.10.22.683864

**Authors:** Haoran Liu, Klaus Benndorf, Yessenbek K. Aldakul, Han Sun

**Affiliations:** Leibniz-Forschungsinstitut für Molekulare Pharmakologie, Berlin 13125, Germany; Institut für Chemie, Technische Universität Berlin, Berlin 10623, Germany; Institut für Physiologie II, Universitätsklinikum Jena, Friedrich-Schiller-Universität Jena, Jena 07740, Germany

**Keywords:** CNG channels, HCN channels, molecular dynamics (MD) simulations, single-channel patch-clamp electrophysiology, selectivity filter

## Abstract

Cyclic nucleotide-gated (CNG) channels and hyperpolarization-activated cyclic nucleotide-gated (HCN) channels are key members of the cyclic nucleotide-activated ion channel family that translate intracellular cyclic nucleotide binding into electrical signals. Functionally, CNG channels drive large inward currents in photoreceptors and olfactory sensory neurons, whereas HCN channels are best known for their roles in pacemaker activity in the heart and the regulation of neuronal excitability. Despite their considerable sequence similarity and conserved overall architecture, these channels exhibit striking differences in ion conductance, K^+^ selectivity, and voltage dependence. Here, we performed microsecond-timescale atomistic molecular dynamics (MD) simulations to directly compare the ion conduction mechanisms of HCN and CNG channels, using the prototypical K^+^-selective channel MthK as a reference. Our simulations reproduced key features observed in single-channel patch-clamp electrophysiology and revealed that distinct selectivity filter architectures and dynamic behaviors are the primary determinants underlying the divergence in ion conductance and K^+^ selectivity between HCN and CNG channels. Together, these results provide a mechanistic framework for understanding the physiological roles of these channels and pave the way for the rational design of cation channels with tailored functional properties.

## 1. Introduction

Cyclic nucleotide-activated ion channels are opened by binding of cyclic nucleotide, thereby converting intracellular signaling into electrical activity.^[1]^ This family comprises cyclic nucleotide-gated (CNG) channels and hyperpolarization-activated cyclic nucleotide-gated (HCN) channels that serve fundamentally different physiological functions, despite their structural similarity.^[1c]^ CNG channels mediate steady inward currents that depolarize photoreceptors and olfactory sensory neurons,^[2]^ whereas HCN channels are essential in generating cardiac pacemaker activity and regulating neuronal excitability.^[3]^ Consistent with these distinct functions, the two channels display strikingly different biophysical properties, particularly in voltage dependence. HCN channels are activated by membrane hyperpolarization, for example after a cardiac action potential, while CNG channels are largely voltage-insensitive. Beyond voltage gating, HCN and CNG channels also differ in ion selectivity and conductance. CNG channels are non-selective cation channels, permeable to both Na^+^ and K^+^, and also allow Ca^2+^ conduction.^[4]^ In contrast, HCN channels exhibit a modest K^+^ selectivity over Na^+^ with a permeability ratio of approximately 4:1.^[5]^ Ion conductance represents another key distinction. CNG channels typically display relatively large unitary single-channel conductance,^[6]^ whereas both homomeric and heteromeric HCN channels exhibit very low conductance, a property that has been the subject of considerable debate in recent electrophysiological and biophysical studies.^[7]^

Cryo-electron microscopy (cryo-EM) structures have provided unprecedented insights into the biophysical properties of both CNG and HCN channels.^[8]^ Although these channels share a broadly similar overall architecture, including the pore domain, voltage-sensing domain (VSD), C-linker, and cyclic nucleotide-binding domain (CNBD) (Figure 1A), they also exhibit pronounced structural differences across these domains. In CNG channels, most positively charged residues within the VSD are positioned such that they are very little affected by transmembrane electric field, providing a plausible explanation for their voltage insensitivity.^[8a]^ This feature sets them apart from other members of the voltage-gated K^+^ channel family, including HCN channels.^[8g,8i]^ Structural comparison of the selectivity filter (SF) highlights further key distinctions. HCN channels share several features with canonical K^+^-selective channels,^[8h]^ particularly at the intracellular segment (around the S3 and S4 ion binding sites), and exhibit an even greater resemblance to the non-selective NaK channel, particularly in the arrangement of the major ion binding sites within the SF.^[9]^ By contrast, the SF of the CNG channels is considerably wider than that of either HCN or K^+^-selective channels, in principle allowing accommodation of partially hydrated cations.^[8a,8c]^ With respect to gating, structural analyses identified a central hydrophobic gate [F389 and V393 in human CNGA1 (hCNGA1)] within the pore domain that controls channel opening in CNG channels (Figure 1B).^[8a,8c,8d]^ In HCN1 and HCN4 channels, however, pore opening is controlled by the coordinated movement of four residues located in the central cavity and lower helix bundles [Y386, V389, T394, and Q398 in human HCN1 (hHCN1) channel; Y507, I511, T515, and Q519 in rabbit HCN4 (rHCN4)].^[8i,8j]^

**Figure 1.**
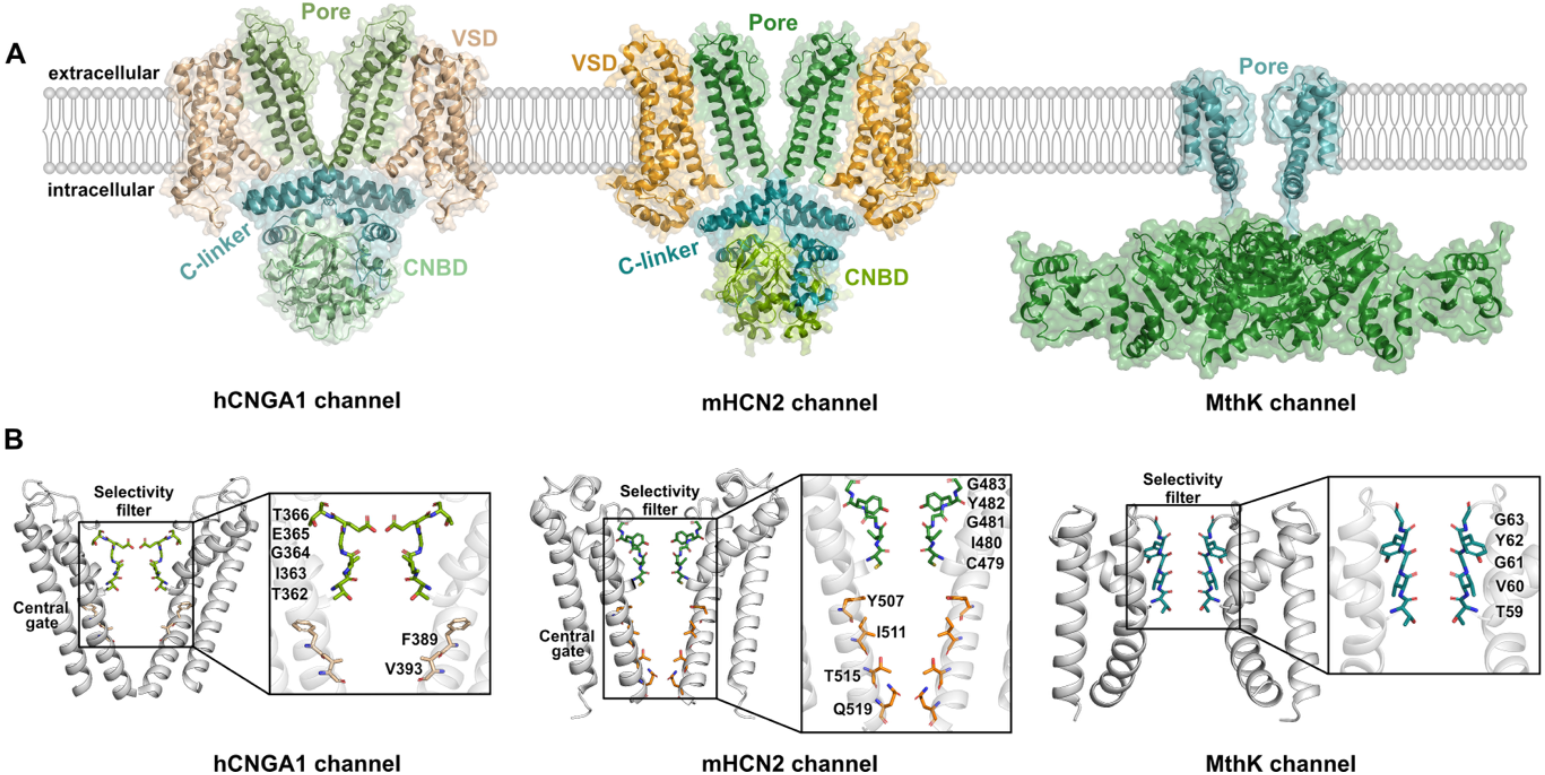
Structural comparison of human CNGA1 (hCNGA1), mouse HCN2 (mHCN2) and bacterial MthK channels. (A) Structures of the (left) open-state hCNGA1 channel (PDB ID: 7LFW^[10]^), (middle) homology model of the mHCN2 channel based on the cryo-EM structure of the open-state of rHCN4 channel (PDB ID: 7NP3^[8j]^), and (right) the X-ray structure of the open-state MthK channel (PDB ID: 6OLY^[11]^). The hCNGA1 and mHCN2 channels are tetramers composed of four subunits, each containing the pore domain (green), voltage-sensor domain (VSD, orange), C-linker (cyan), and cyclic nucleotide-binding domain (CNBD, light green). In MthK, the transmembrane pore domain (cyan) is linked to the cytoplasmic regulator of conductance of K^+^ (RCK) domain (green), while only the pore domain was included in the simulations. For clarity, only two diagonally opposed subunits are shown in all structures. (B) Structural details of the pore domains in the open-state for the hCNGA1, mHCN2, and MthK channels. Key residues in the SF and central gate are highlighted and labeled.

Building on the availability of cryo-EM structures of both HCN and CNG channels, a number of atomistic molecular dynamics (MD) simulations have been performed to investigate the dynamic mechanisms underlying ion permeation, gating and ligand binding. Simulations of rHCN4 channels revealed cation-specific differences in ion binding affinity and SF plasticity, consistent with its weak cation selectivity.^[12]^ Similarly, MD simulations of the HCN1 channel uncovered distinct ion binding profiles in the SF compared to voltage-gated K^+^ (K_v_) channels, accompanied by increased pore flexibility.^[13]^ More recently, we demonstrated that differences in ion occupancy at the extracellular side of mHCN isoforms contribute to variations in single-channel conductance. In particular, negatively charged residues at the extracellular entrance of the SF enhances ion binding and thereby promote multi-ion knock-on conduction. In addition, advanced enhanced sampling simulations of hHCN1 channels proposed an atomistic coupling mechanism between the voltage-sensing domain (VSD) and the pore domain, and further suggested a modulatory role of lipid in hyperpolarization-dependent gating.^[14]^ Regarding ligand binding, mass spectrometry analyses combined with MD simulations identified a distinct binding pocket present only in the resting conformation of the voltage sensor.^[15]^ For the hCNGA1 channel, our recent work showed that the unusually wide SF accommodates both K^+^ and Na^+^ in a partially hydrated state, facilitated by a flexible SF and a dynamic gate region.^[16]^

In the present study, we systematically compared the ion conductance, hydration states, and the free energy profiles of K^+^ permeation in mHCN2 and hCNGA1 channels using atomistic MD simulations performed under comparable conditions. The simulated conductance of both channels reproduced key features observed in single-channel recordings of both channels under similar conditions. Using the K^+^-selective MthK channel as an additional reference, we demonstrated that differences in the size and dynamic behavior of the SF underlies the distinct conductance and selectivity characteristics of HCN and CNG channels.

## 2. Results

### 2.1. Simulated conductance of mHCN2 and hCNGA1 channels reproduced key single-channel conductance properties

We recently reported atomistic MD simulations of ion permeation in mHCN1-4 and hCNGA1 channels.^[7b]^ These previous mHCN1-4 simulations were carried out using the AMBER19SB force field to allow direct comparison with earlier MD studies of the related rHCN4 channel by Hamacher *et al*.^[12]^. In contrast, simulations of hCNGA1 employed the CHARMM36m force field, as the AMBER force field produced unrealistically low Na^+^ conductance for the non-selective cation channels^[16]^. To enable a more direct comparison of conductance properties between mHCN and hCNGA1 channels, we have performed additional MD simulations of the full-length mHCN2 channel using the CHARMM36m force field. Among the four isoforms of mHCN channels, mHCN2 exhibits the largest single-channel conductance and was therefore chosen for comparative analyses with hCNGA1. As in our earlier work, ion conductance was estimated from five independent 1 μs simulations. For hCNGA1, simulations were performed at physiologically relevant voltages (+100 mV and −100 mV), whereas substantially higher transmembrane voltage (−700 mV) was required for mHCN2 to obtain statistically meaningful conductance estimates, owing to its intrinsically low ion conductance that was shown in our previous study^[7b]^.

In this study, we also performed single-channel patch-clamp recordings of both mHCN2 and hCNGA1 channels in *Xenopus* oocytes under comparable conditions to enable direct comparison with simulations. Representative traces and current-amplitude diagrams revealed distinct conductance properties of hCNGA1 channels at positive versus negative transmembrane voltages, as well as clear differences between hCNGA1 and mHCN2 channels (Figure 2A/B): Consistent with previous reports, hCNGA1 channels exhibited an approximately 22-fold higher single-channel conductance than mHCN2 channels (−100 mV versus −130 mV, respectively) (Figure 2C). In addition, the single-channel conductance of hCNGA1 channels at −100 mV is considerably larger than at +100 mV and the width of the distribution is clearly larger due to a flicker.

**Figure 2.**
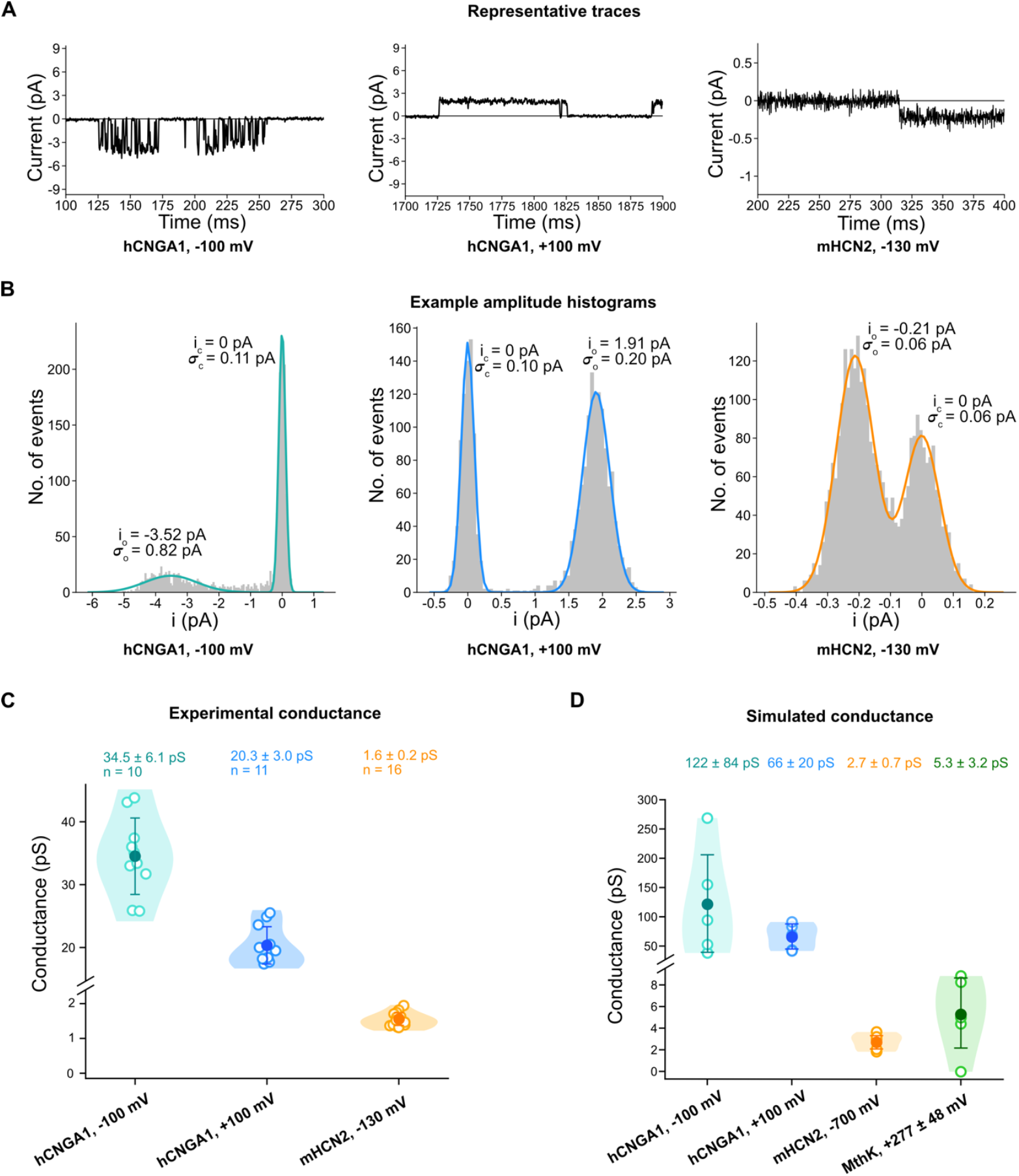
Comparison of the experimental and simulated conductances of hCNGA1 and mHCN2 channels. (A) Representative single-channel current traces for hCNGA1 channels at −100 mV and +100 mV as well as of mHCN2 channels at −130 mV. (B) Amplitude histograms of the traces in A. The baseline current peak (i_c_) was shifted to zero. The histograms were fitted by the sum of two Gaussian functions (colored curves) yielding the indicated single-channel current i_o_ and the corresponding standard deviations *σ*_o_ and *σ*_c_. (C) Conductance values for the hCNGA1 channels at −100 mV and +100 mV as well as of mHCN2 channels at −130 mV. Open circle represents the experimental conductance from one individual single-channel recording, while the closed circle shows the average conductance from several measurements. The error bar represents the standard deviation. (D) Conductance values derived from simulations of an hCNGA1 channel (cyan, −100 mV; blue, +100 mV), mHCN2 channel (orange) and bacterial MthK channel (green; pore domain only). Open circle represents the simulated conductance from one individual simulation run, while the closed circle shows the average conductance from five parallel simulation runs. The error bar represents the standard deviation.

Comparison of simulated and experimental conductance for hCNGA1 and mHCN2 channels showed that the simulations qualitatively reproduced the trend of a much higher unitary conductance in hCNGA1 relative to mHCN2. For mHCN2, the simulated conductance of 2.7±0.7 pS closely matched the experimental values of 1.6±0.2 pS (Figure 2C/D). In contrast, the simulated K^+^ conductance of hCNGA1 at both positive and negative voltages was approximately 3.5-fold higher than the corresponding experimental measurements (Figure 2C/D). Notably, the simulations also captured two key experimental features of the hCNGA1 channel: (i) a higher inward than outward conductance, and (ii) a broader distribution of single-channel conductance levels at the negative voltages.

We recently reported atomistic MD simulations of K^+^ permeation through the MthK channel using a computational protocol similar to that applied here for hCNGA1 and mHCN2 channels.^[17]^ The simulated K^+^ outward conductance of MthK was 5.3±3.2 pS (Figure 2D), which lies between the values obtained from hCNGA1 and mHCN2 simulations. This simulated conductance is approximately 10-fold lower than that measured previously in single-channel recordings of purified MthK reconstituted in lipid bilayer.^[18]^ A likely explanation for this discrepancy is that only the transmembrane domain of MthK was included in our simulations, during which partial closure of the lower helix bundle was observed. As we were unable to measure the single-channel conductance of MthK in *Xenopus* oocytes, these simulation data are not intended for direct quantitative comparison with experimental conductance values. Instead, they serve as an additional reference to aid in understanding the conductance and selectivity differences between the two cyclic nucleotide-gated channels discussed in the following sections.

### 2.2. Comparison of the pore radius and hydration level of the pore in mHCN2, hCNGA1, and MthK channels

To gain a deeper understanding of the distinct conductance differences observed in the simulations, we compared the pore radii of the mHCN2, hCNGA1, and MthK conformations sampled during the MD simulations (Figure 3). Pore radii and profiles were calculated using the program HOLE^[19]^. As shown in Figure 3, the SF of hCNGA1 widened considerably during the simulations, while the gate region sampled conformations intermediate between open and closed states. In contrast, the pore domain of mHCN2 channel remained largely stable throughout the simulations. The overall gate dimensions sampled by hCNGA1 and mHCN2 were comparable, indicating that the gate size is not the primary determinant of their markedly different conductance properties. Instead, the SF constitutes the narrowest region of the pore in both channels, with hCNGA1 exhibiting a substantially wider SF than mHCN2. In the case of MthK, the SF remained stable and also formed the narrowest constriction within the pore. The gate shifted slightly towards a more closed conformation, likely because only the transmembrane pore domain was included in the simulations, which led to some instability of the gate. Nevertheless, the pore remained sufficiently open to permit K^+^ conduction.

**Figure 3.**
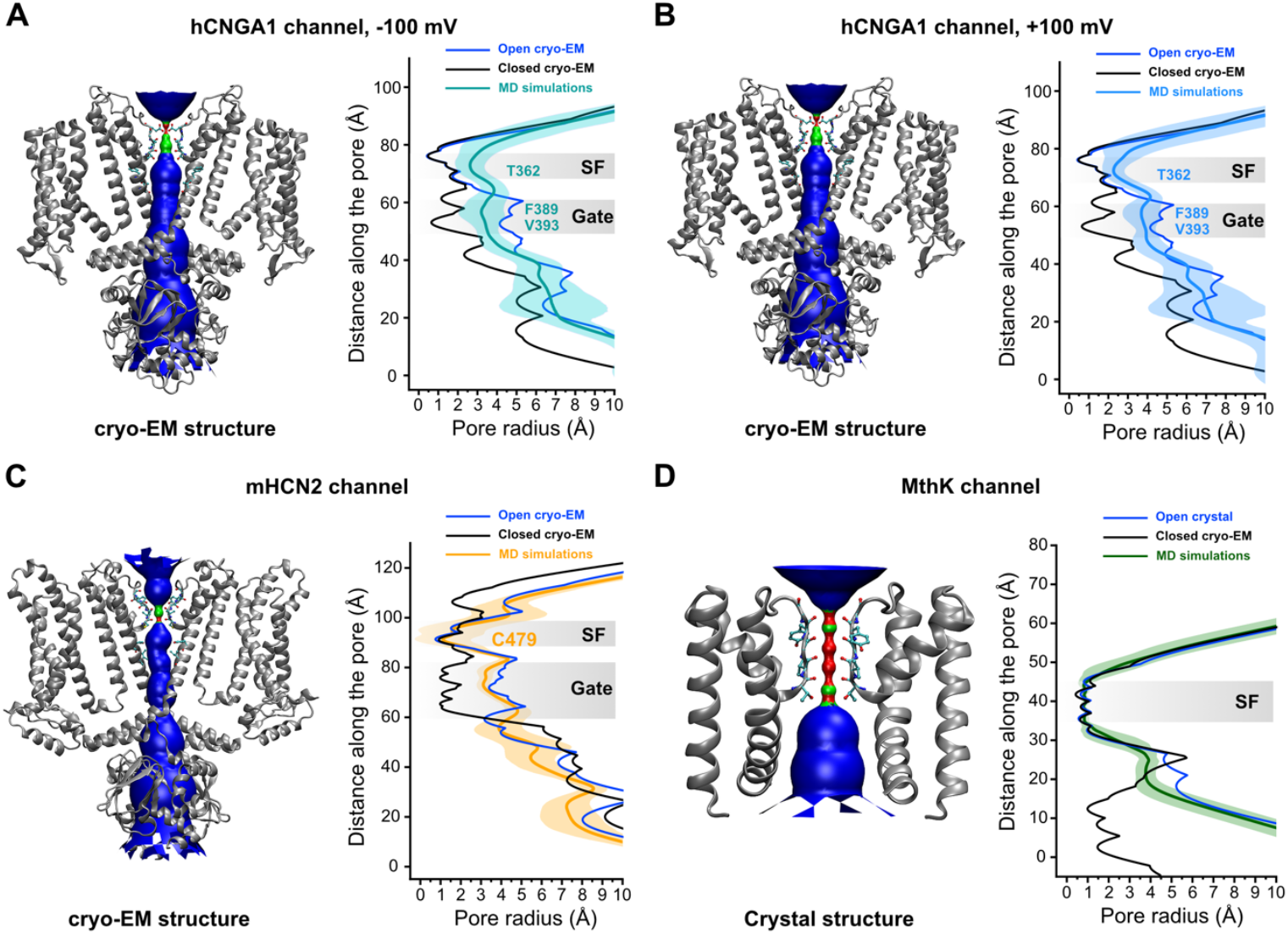
Pore radius profiles along the central axis during MD simulations. Analysis of simulations of (A) hCNGA1 channel at −100 mV, (B) hCNGA1 channel at +100 mV, (C) mHCN2 channel at −700mV, and (D) MthK channel at +277±48 mV. (left) HOLE pore profile of the open conformation in the original cryo-EM or crystal structures; (right) Comparison of pore radius profiles along the central axis between open and closed cryo-EM or X-ray structures, together with those derived from MD simulations. The dark-colored line represents the average pore radius, and the light-colored shading indicates the standard deviation calculated from 10,000 snapshots across five parallel simulation runs. The SF and gate regions are highlighted in gray, with key residues indicated.

To further assess hydration properties of the central cavity, we calculated the average number of water molecules within the pore across individual simulations (Figure 4). Among the three channels, hCNGA1 channel displayed the highest average hydration level, whereas MthK showed the lowest. Notably, the hydration levels of the hCNGA1 channel were similar under positive and negative voltages, although the distribution was substantially broader at negative voltage, consistent with the greater variability in conductance observed across individual runs. However, comparison across all three channels revealed no clear correlation between pore hydration and simulated conductance. Therefore, pore hydration alone does not appear to be the determining factor underlying the conductance differences observed among these channels.

**Figure 4.**
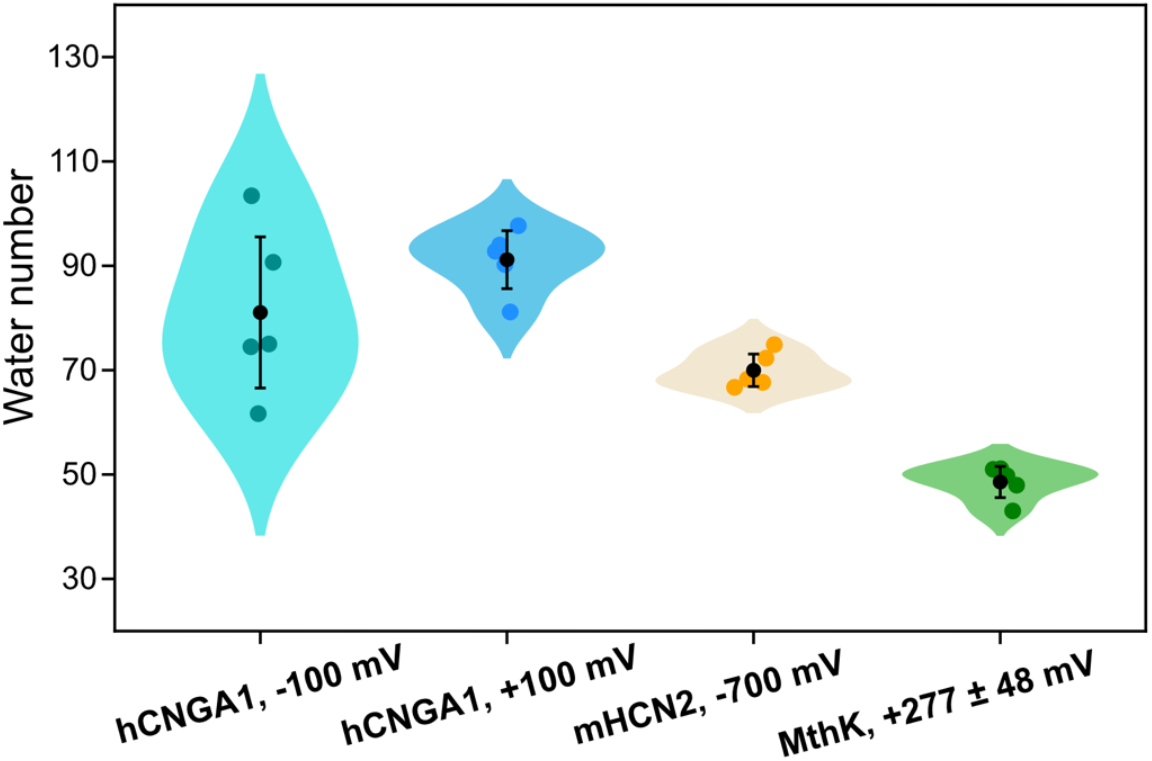
Number of averaged water molecules in the pore cavity of the hCNGA1, mHCN2, and MthK channels. Each colored round dot represents the average number of water molecules calculated from 20,000 snapshots in one individual simulation run. The black dot represents the average value from five independent simulation runs and the error bar shows the standard deviation.

### 2.3. Comparison of the free-energy profiles of K^+^ permeation

Next, we compared the free-energy profiles of K^+^ permeation derived from simulations for hCNGA1, mHCN2, and MthK channels (Figure 5). In hCNGA1, the SF region exhibited the most negative ΔG values, clearly indicating that this region serves as the primary ion binding region of the channel. Owing to the relatively large dimensions of the filter, ions can move with minimal restriction, resulting in the absence of a distinct energy barrier for ion permeation within the filter. In contrast, the mHCN2 channel displayed two major K^+^ binding sites within the SF, consistent with the S3 and S4 binding sites identified in cryo-EM structure. These two binding sites are separated by an energy barrier of about 5 kcal/mol. For MthK, five distinct ion binding sites were observed, with energy barriers of similar magnitude to those observed in mHCN2. In summary, comparison of the K^+^ permeation free-energy profiles across the three channels reveals that the wide SF of hCNGA1 presents a relatively low energy barrier due to its wide dimensions, whereas distinct binding sites and comparatively high energy barriers characterize the mHCN2 and MthK channels.

**Figure 5.**
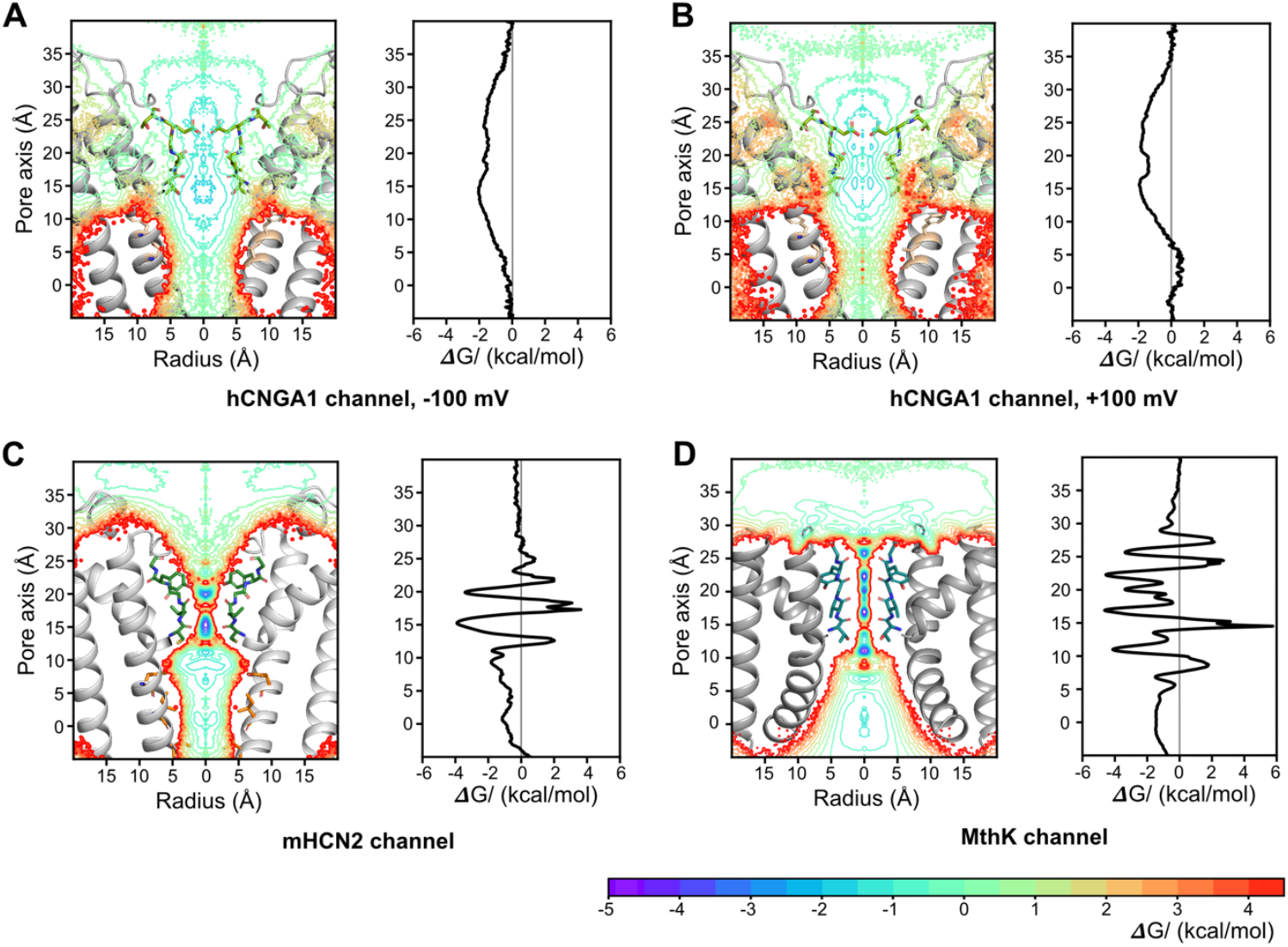
Free energy profiles of ion conduction in (A) hCNGA1 channel at −100 mV, (B) hCNGA1 channel at +100 mV, (C) mHCN2 channel at −700mV, and (D) MthK channel at +277±48 mV derived from MD simulations. (left) Two-dimensional free energy profiles along the pore during ion permeation; (right) One-dimensional free energy profiles along the pore during ion permeation. Free energy profiles are derived from the ion occupancy along the pore, where the ion occupancy in number of ions per 0.001 Å^3^ per 50 ps was normalized according to the volume change along the radius.

Additionally, we observed that the ΔG values in the pore cavity are less negative in hCNGA1 than in mHCN2 and MthK channels, likely due to the hydrophobic nature of the cavity environment. Overall, the free-energy profile of the mHCN2 channel more closely resembles that of MthK than of hCNGA1, despite the strong structural similarity and sequence conservation between mHCN2 and hCNGA1 channels.

### 2.4. Comparison of the SF dynamics

A previous MD study suggested that ion-induced structural plasticity of the SF underlies the weak K^+^ selectivity of the HCN4 channel[12b]. Here, we revisited this mechanism by comparing the SF flexibility of hCNGA1, mHCN2, and MthK channels, which exhibit distinct degree of K^+^ selectivity: hCNGA1 is completely non-selective, MthK is highly selective, and mHCN2 displays weak K^+^ selectivity. To this end, we calculated residue-wise root-mean-square fluctuation (RMSF) values for the SF residues from MD simulations of all three channels. As shown in Figure 6 and Figure S1, the RMSF values followed the same trend as their selectivity: the highest RMSF values, reaching up to 0.3 nm, were obtained for hCNGA1 channel, the lowest values of around 0.05 nm for MthK, and intermediate values for mHCN2. The standard deviations across individual simulation runs followed the same pattern. RMSF values for hHCNA1 simulations at positive and negative voltages were highly comparable and remained the highest among three investigated channels. Interestingly, both hCNGA1 and mHCN2 channels exhibited higher flexibility at the extracellular side of the filter compared to the intracellular side. Overall, the pronounced differences in SF flexibility among the three channels support previous findings that conformational plasticity of the SF is an important determinant for cation non-selectivity.

**Figure 6.**
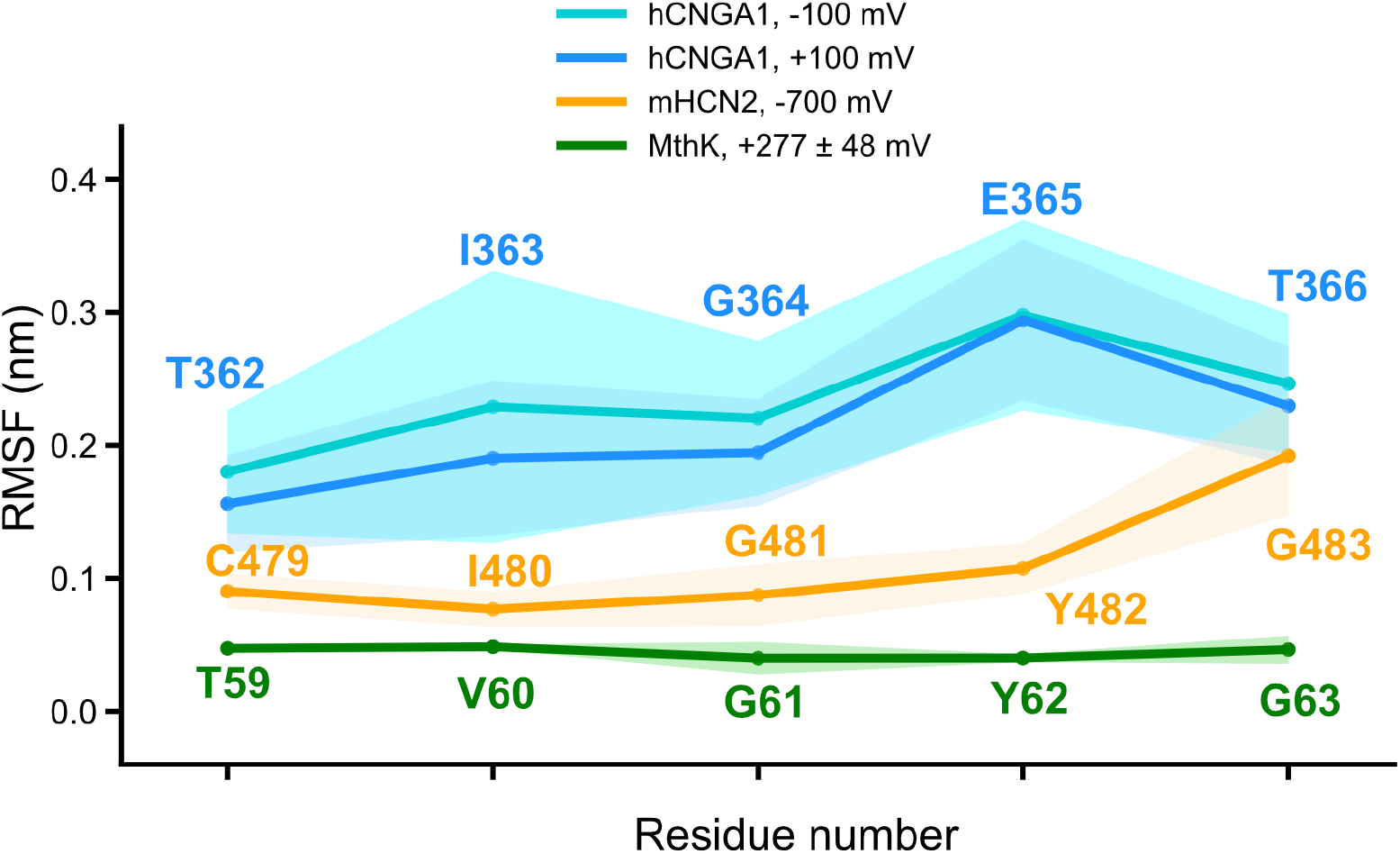
Root-mean-square-fluctuation (RMSF) of the SF from MD simulations of hCNGA1, mHCN2, and MthK channels. For each simulation setup, the deep-colored line represents the mean RMSF across five independent runs, while the light-colored shading indicates the standard deviation.

### 2.5. Comparison of the hydration states of K^+^ in the SF during their permeation

Several previous studies have emphasized the importance of ion dehydration within the SF for achieving high K^+^ selectivity. To assess whether this mechanism also applies to the three channels investigated here, we calculated the number of coordinating water molecules and protein carbonyl oxygens within the first hydration shell of K^+^ ions as they traversed the pore during the simulations (Figure 7). In bulk solution, K^+^ ions are coordinated on average by seven molecules in all channels. Within the SF region, however, distinct differences emerged. In the K^+^-selective MthK channel, K^+^ ions become fully dehydrated at several major binding sites within the SF. By contrast, in the non-selective hCNGA1 channel, K^+^ ions lose only about two water molecules, which are replaced by coordination with the carbonyl oxygens of SF residues at the primary binding site S4. The mHCN2 channel again displayed intermediate behavior: at its primary S4 site, carbonyl coordination competes with water, leaving roughly only two water molecules in the hydration shell of K^+^ ions. Taken together, these results demonstrate that the extend of K^+^ dehydration with the SF correlates closely with the degree of K^+^ selectivity across the three channels: complete dehydration in MthK corresponds to strong K^+^ selectivity, partial dehydration in mHCN2 to weak selectivity, and minimal dehydration in hCNGA1 to non-selectivity of monovalent cations.

**Figure 7.**
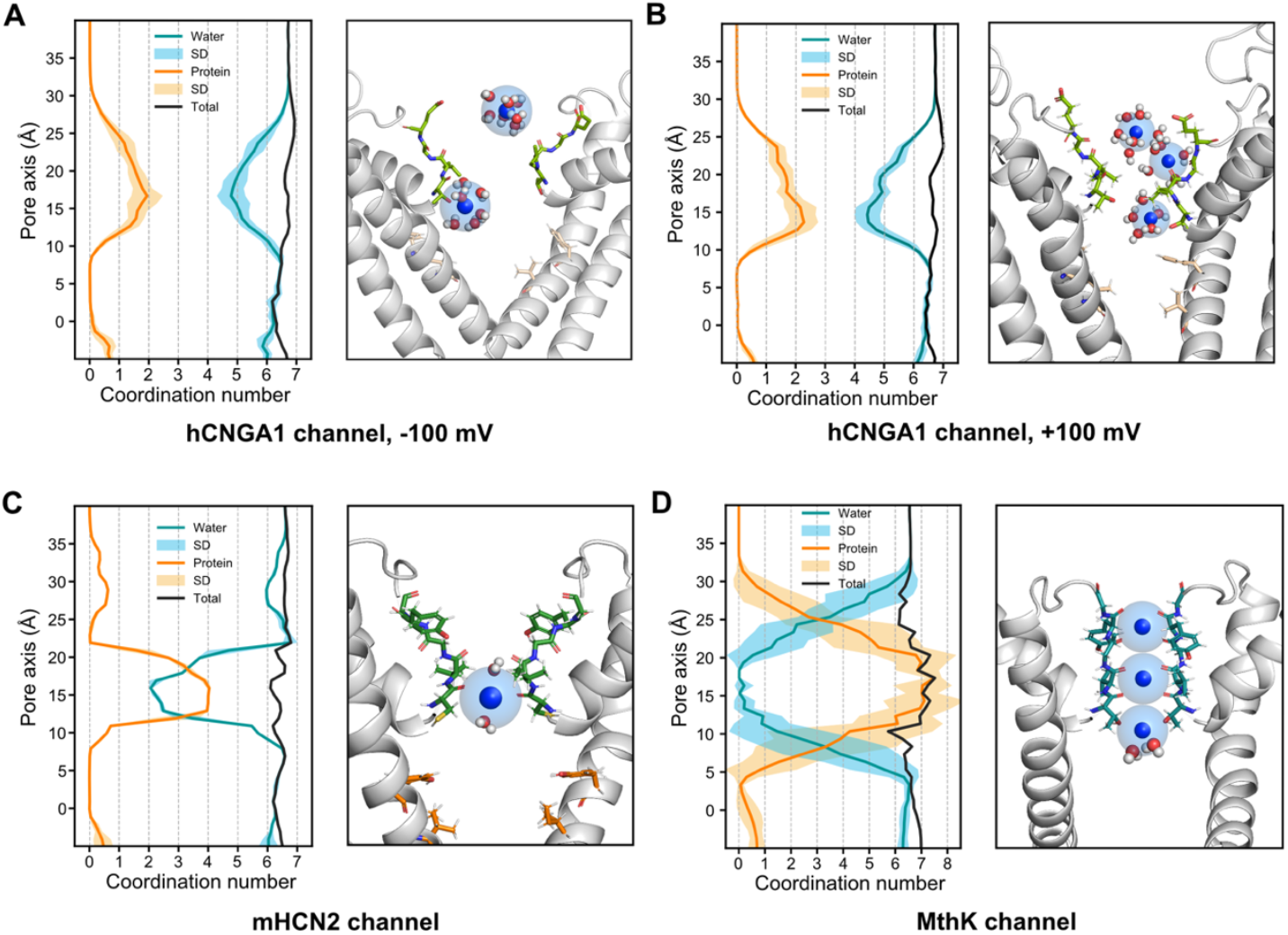
Hydration profiles of the (A) hCNGA1 c(hannel at −100 mV, (B) hCNGA1 channel at +100 mV, (C) mHCN2 channel at −700mV, and D) MthK channel at +277±48 mV derived from MD simulations. (left) The orange line represents the average number of protein carbonyl oxygens within the first hydration shell of the K^+^, and the light orange shading indicates the standard deviation from five independent simulation runs. The blue line represents the average water oxygens within the first hydration shell, and the light blue shading indicates the standard deviation from five independent simulation runs. The black line represents the total number of coordinating oxygens within the first hydration shell of the K^+^ ion. (right) Representative simulation snapshots of K^+^ ions (blue) are shown, with the first hydration shell depicted as a transparent sphere.

## 3. Discussion

HCN and CNG channels are members of the voltage-gated K^+^ channel family and are activated by the binding of cyclic nucleotides. Despite their similar overall architecture, they exhibit pronounced differences in ion conductance, selectivity and voltage sensitivity, which account for their different physiological functions. In this work, we focused on elucidating the mechanistic origins of the differences in ion conductance and selectivity between HCN and CNG channels. Our main conclusion was drawn based on a comparison between conductance properties obtained from atomistic MD simulations under transmembrane voltages and single-channel patch clamp electrophysiology data collected under comparable conditions, combined with a systematic analysis of the simulation results. Furthermore, we employed the bacterial K^+^-selective channel MthK as a reference in our simulations. This comparison proved particularly valuable for clarifying the structural and energetic determinants underlying the differing degrees of K^+^ selectivity observed in HCN and CNG channels.

Experimentally, we observed that the hCNGA1 channel exhibited more than 20-fold higher conductance than the mHCN2 one in single-channel recordings measured under closely matched conditions. In the simulations, the conductance difference appeared even larger, reaching nearly 45-fold between hCNGA1 and mHCN2 channels. We have recently discussed that one possible reason for the much higher simulated conductance of hCNGA1 channel than experimentally observed values could be that classical additive force field parameters may not optimally capture the transient and complex interactions among ions, water, and protein residues within the relatively dynamic and wide SF.^[17]^ Nevertheless, our simulations qualitatively reproduced three key conductance properties of the hCNGA1 and mHCN2 channels from the single-channel electrophysiology experiments: (i) a markedly higher inward K^+^ conductance compared to outward conductance; (ii) a generally broader conductance distribution for hCNGA1 than for mHCN2; (iii) a broader conductance distribution for hCNGA1 at negative voltages (inward flow) than at positive voltages (outward flow).

For the higher inward K^+^ conductance compared to the outward conductance in the hCNGA1 channel, we propose that the asymmetric distribution of charge and hydrophobicity in the pore may be one of the underlying reasons (Supplementary Figure S2). Due to the relatively high negative electrostatic potential, a number of K^+^ ions accumulate at the extracellular side, where a multi-ion knock-on conduction mechanism may enhance the rate of ion flow. In contrast, the intracellular vestibule is strongly hydrophobic, resulting in very few ions accumulating on the intracellular side. A similar mechanism has been proposed by Hilder *et al*. to explain similar behavior in inwardly rectifying K^+^ channels using Brownian dynamics simulations.^[20]^

Next, we turned our focus to the broader conductance distribution observed for the hCNGA1 channel compared to mHCN2, as well as the broader distribution for hCNGA1 at negative voltages relative to positive ones. Analysis of pore profiles from the MD simulations revealed that the pore, and in particular the gate residues of hCNGA1, is substantially more flexible than in mHCN2. This increased flexibility results in a more dynamic hydration level within the pore cavity region below the SF. As we have previously suggested, hydration level in the gate region often correlates with conductance in the simulations.^[16]^ We therefore propose that the dynamic pore geometry and fluctuating hydration state in the cavity of hCNGA1 channel underlie the broad single-channel conductance distribution observed experimentally. Consistent with the smaller variation in conductance distribution at positive voltages compared to negative ones, we also observed a narrower distribution of cavity hydration under positive voltages.

To understand the remarkable difference in conductance between hCNGA1 and mHCN2 channels, we first compared their gate distances, which remained rather similar throughout the simulations. Likewise, the hydration state of the central cavity did not differ remarkably, indicating that pore dehydration also cannot account for the conductance difference. Notably, the most pronounced difference emerged in the SF. The cryo-EM structure of hCNGA1 channel already revealed a substantially wider SF compared to mHCN2, and this difference became even more pronounced during the simulations, accompanied by enhanced dynamics. Free-energy profiles of ion conduction provided additional insights. In hCNGA1, the large SF dimension does not impose a high energy barrier for cation conduction. By contrast, both mHCN2 and MthK channels exhibit well-defined ion binding sites within the SF, with MthK showing an even greater number. These binding sites are separated by energy barriers of up to 5 kcal/mol. A previous simulation work of HCN4 also suggested that the high energy barrier within the SF accounts for the low unitary conductance of the HCN4 channel.^[12a]^ We therefore propose that the pronounced difference in SF dimension, and the resulting disparity in energy barriers, largely accounts for the substantial difference in ion conductance between hCNGA1 and mHCN2 channels.

In addition, analyses of the SF dynamics of the three investigated channels, reflected by the RMSF profiles, revealed a strong correction between K^+^ selectivity and SF plasticity. The K^+^-selective MthK channel exhibits a highly rigid SF, whereas the non-selective hCNGA1 shows pronounced flexibility in the filter, with the weakly K^+^-selective mHCN2 lying in between. Previous simulations of the HCN4 channel likewise highlighted the importance of SF flexibility in mediating weak K^+^ selectivity. ^[12]^ For the NaK channel, we have proposed previously that conformational plasticity in the SF of both wild-type and mutant channels underlies their cation non-selectivity.^[21]^ Notably, NaK channel has long been considered as a bacterial homologue of CNG and HCN channels. Taken together, these findings collectively suggest that cation channels tune their K^+^ selectivity through the degree of SF flexibility. A more flexible and wider SF can accommodate hydrated Na^+^ ions, whereas K^+^ ions, which have a lower dehydration energy, can permeate in both hydrated and dehydrated forms. Because Na^+^ dehydration is energetically much more unfavorable than that of K^+[22]^, increased SF flexibility reduces the selectivity barrier. Consistent with this proposal, our comparison of K^+^ hydration levels in the SF across the three channels shows a direct correlation between hydration, SF flexibility, and the degree of K^+^ selectivity.

In conclusion, we carried out a systematic investigation of the mechanisms underlying the differences in ion conductance and selectivity between CNG and HCN channels. Building on our recent advances in single-channel patch-clamp recordings at low-noise conditions^[16]^, we obtained high-quality single-channel electrophysiology data for hCNGA1 and mHCN2 channels measured under comparable experimental conditions. Complementary atomistic MD simulations on microsecond timescale reproduced the key features and trends observed in the single-channel recording. The simulations revealed that a wider filter dimension combined with multi-ion knock-on conduction supports high conductance, whereas a narrower and conformationally more rigid filter that enforces K^+^ dehydration governs strong K^+^ selectivity.

## 4. Methods

### 4.1. Atomistic molecular dynamics simulations

MD simulations started from the high-resolution cryo-EM structure of the homomeric hCNGA1 channel in the open state (PDB ID: 7LFW). The starting structure of mHCN2 channel was generated from the homology modelling of the open conformation of the rabbit HCN4 channel (PDB ID: 7NP3)^[8j]^. The MD simulations of MthK channel started from the open conformation of crystal structure (PDB ID: 3LDC)^[23]^ and have been reported previously^[17]^. The simulations of CNGA1 channel and mHCN2 channel were all starting from the full-length constructs, while the simulations of MthK channel contained only the isolated transmembrane pore domain, a strategy frequently employed in MD simulations of ion channels to mitigate computational complexity^[14,24]^. To prevent any artifacts originating from charges at the N- and C-termini, the structures were neutralized by adding acetyl and N-methyl caps using PyMOL^[25]^ Builder option.

All simulations were performed using the CHARMM36m force field^[26]^ for a better comparison. All channel structures were embedded within a palmitoyloleoyl phosphatidylcholine (POPC) lipid bilayer with TIP3P (Transferable Intermolecular Potential with Three Points) water model^[27]^ using CHARMM-GUI^[28]^ webserver in all simulations. The KCl concentration was set to 150 mM in hCNGA1 simulations, 900 mM in mHCN2, and 600 mM in MthK simulations, respectively (Supplementary Table S1).

After the simulation systems were embedded into a POPC lipid bilayer with ions and water in a box, the simulation systems were energy minimized and equilibrated. The energy minimization was done to reduce the system maximum force to below 1000 kJ/mol/nm in 50,000 steps. After the energy minimization, an equilibration in six steps was performed using default scripts provided by the CHARMM-GUI. For mHCN2 simulations, an extra 50 ns equilibration without any restraint was performed.

For simulations of the hCNGA1 channel and mHCN2 channel, the transmembrane electric potential was directly generated by an external electric field applied along the z-axis (pore axis) (Supplementary Table S1). The voltage was calculated with:

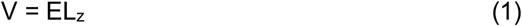

where E is the applied electric field and L_z_ is the length of the simulation box along the z-axis[29]. For MthK channel simulations, computational electrophysiology (CompEL) setup was used (Supplementary Table S1)^[30]^. To establish a transmembrane potential, a copy of the equilibrated system was stacked along the pore axis, generating two compartments (an inner and an outer). After building the two-bilayers system we conducted another round of equilibration for 20 ns. For the production run, a charge imbalance of 2 *e* between the two compartments separated by two lipid bilayers resulted in +277 ± 48 mV and −277 ± 48 mV, respectively. During the MD simulations, ions passing through the pore were monitored, and the charge imbalance was maintained by exchanging the same species of ion in one compartment with a water molecule from the other compartment^[30]^. The resulting transmembrane voltage can be calculated by double integration of the charge distribution using the Poisson equation as implemented in the GROMACS tool *gmx potential*^[31]^.

All atomistic MD simulations were run with GROMACS software package (version 2021.2 and 2023.3)^[32]^. Short-range electrostatic interactions were calculated with a cutoff of 1.0 nm, whereas the long-range electrostatic interactions were treated using the particle mesh Ewald method^[33]^. The cutoff for van der Waals interaction was set to 1.0 nm. The simulations of hCNGA1 channel and mHCN2 channel were performed at 300 K with an enhanced Berendsen thermostat (GROMACS V-rescale thermostat^[34]^ while the simulation of MthK channel was set to 303.15K. The Parrinello-Rahman barostat^[35]^ was employed to keep the pressure within the system remaining at 1 bar. All bonds were constrained with the Linear Constraint Solver (LINCS) algorithm^[36]^. The protonation states of all titratable residues were assigned according to the standard protonation states at pH 7. All simulations were performed with an integration time step of 2 fs. The root-mean-square-deviation (RMSD) of the MD simulations of all systems are shown in Supplementary Figure S3.

All trajectories were analyzed with GROMACS toolkits and Python3 using MDAnalysis^[37]^, Numpy^[38]^, Matplotlib^[39]^, Pandas^[40]^ and SciPy^[41]^ packages. We calculated the pore radius of the channels using HOLE program^[19]^. Pore radius profiles during the simulations were computed from 10,000 frames, corresponding to 2,000 frames extracted from a representative trajectory out of five independent MD simulation replicates. Ion permeation events were calculated when a cation permeated through the entire pore from the SF to the gate. To determine the ion hydration states during permeation, we calculated the number of ion-coordinating oxygens from both water molecules and protein carbonyls within each ion’s first solvation shell. Hydration shells are dynamic; here, we defined the waters of hydration using the radii corresponding to the minimum in the radius of gyration profiles: 3.1 Å for Na^+^ and 3.4 Å for K^+^.^[42]^ As for the free energy profile, we calculated it based on the concentration of ions in the pore by the Gibbs free energy change for diffusion or passive transport:

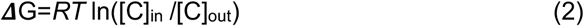

where R = *N*_*A*_ *\* k*_*B*_ ≈ 8.314 J/(mol*K), *N*_*A*_ is the Avogadro constant and *k*_*B*_ is the Boltzmann constant. The free energy was the average value from 5 independent simulation runs. Molecular visualizations were made with PyMol and Visual Molecular Dynamics (VMD)^[43]^.

All simulation details were summarized in Supplementary Table S1 and S2.

### 4.2. Patch-clamp measurements

Oocytes were surgically harvested under anesthesia (0.3% 3-aminobenzoic acid ethyl ester) from adult females of *Xenopus laevis*. The procedures regarding the *X. laevis* frogs were approved by the animal ethics committee of the Friedrich Schiller University Jena (UKJ-18-008 from 09 May 2018). The respective protocols were performed in accordance with the approved guidelines. Extreme efforts were made to reduce the stress and to keep the number of frogs to a minimum.

Single-channel currents were recorded with the patch-clamp technique at low-noise conditions^[16]^ from patches of *Xenopus* oocytes expressing the respective channels. Ooycyte preparation, cRNA injection followed the same protocol as previously reported^[16]^. A K^+^ rich bath solution (in mM: 100 KCl, 10 EGTA, 1 MgCl_2_, 10 Hepes (pH 7.2) zeroed the resting potential of the oocytes. All recordings were performed at room temperature (20-22 ^o^C). The patch pipettes were fabricated from quartz tubing (outer and inner diameter 1.0 and 0.5 mm, respectively) using the P-2000 puller, Sutter Instruments, Novato (CA), USA (Science Products GmbH, Hofheim, Germany) to keep the noise low [44]. The resistance of the pipettes was 7 to 14 MΩ. The pipette solution contained 120 KCl, 10 Hepes, 1 MgCl_2_, 1 CaCl_2_ (pH 7.2) respectively.

hCNGA1 channels were measured in the inside-out configuration with 46.5 μM cGMP in the bath, a concentration at the EC_50_ value^[45]^. The sensitivity of the hCNGA1 channels to cGMP was tested. mHCN2 channels were measured in the cell-attached configuration. They were identified by the slow voltage-dependent activation^[16]^.

Currents were recorded with an Axopatch 200B amplifier (Axon Instruments Inc., Foster City (CA), USA) in the capacitive mode. Stimulation and data recording were performed with the ISO3 hard- and software (MFK, Niedernhausen, Germany). The sampling rate was 10 kHz. The on-line filter of the amplifier (4-pole Bessel) was set to 1 kHz. Reasonable patches had a seal resistance of >100 GΩ.

## Data availability

All data supporting the findings of this study are included in this manuscript and in the Supplementary Information. All source data, Supplementary Movies and the simulation input files, comprising the starting configurations and all necessary parameters for performing the MD simulations are deposited in Zenodo under accession code https://doi.org/10.5281/zenodo.17379854

## Code availability

MD simulations data were generated using GROMACS 2021.1/2023.3. All trajectories were analyzed with GROMACS tools and Python using MDAnalysis, Numpy, Matplotlib, Pandas, and SciPy.

## Acknowledgements

This work was funded by the Leibniz-Forschungsinstitut für Molekulare Pharmakologie (FMP) and Deutsche Forschungsgemeinschaft (DFG) CRC1078 ‘Protonation Dynamics in Protein Function’, RU2518 DynIon, and a Leibniz Collaborative Excellence project (to H.S.). The MD simulations were performed with resources provided by the North-German Supercomputing Alliance (HLRN), the high-performance computer “Lise” at the NHR Center (NHR@ZIB) and Erlangen National High Performance Computing Center (NHR@FAU). We are grateful to Uta Enke, Claudia Ranke and Ulrike Singer for excellent technical support. We also acknowledge the technical support from Dr. Songhwan Hwang and Dr. Tillmann Utesch.

## Author contributions

H.S. and H.L. designed and directed the project. H.L. and Y.A. performed the MD simulations. K.B. performed the single-channel patch-clamp recordings. H.L. analyzed the MD simulations data with the contributions from Y.A. The manuscript was written by H.L., K.B and H.S.

## References

[1] a) U. B. Kaupp, R. Seifert, Physiol Rev 2002, 82 (3), 769, 10.1152/physrev.00008.2002; b) M. Biel, S. Michalakis, Handb Exp Pharmacol 2009, (191), 111, 10.1007/978-3-540-68964-5_7; c) K. B. Craven, W. N. Zagotta, Annu Rev Physiol 2006, 68, 375, 10.1146/annurev.physiol.68.040104.134728; d) M. Biel, C. Wahl-Schott, S. Michalakis, X. Zong, Physiol Rev 2009, 89 (3), 847, 10.1152/physrev.00029.2008.

[2] a) E. E. Fesenko, S. S. Kolesnikov, A. L. Lyubarsky, Nature 1985, 313 (6000), 310, 10.1038/313310a0; b) T. Nakamura, G. H. Gold, Nature 1987, 325 (6103), 442, 10.1038/325442a0.

[3] a) D. DiFrancesco, Nature 1986, 324 (6096), 470, 10.1038/324470a0; b) C. S. Chan, R. Shigemoto, J. N. Mercer, D. J. Surmeier, J Neurosci 2004, 24 (44), 9921, 10.1523/JNEUROSCI.2162-04.2004.

[4] S. Frings, J. W. Lynch, B. Lindemann, J Gen Physiol 1992, 100 (1), 45, 10.1085/jgp.100.1.45.

[5] a) R. B. Robinson, S. A. Siegelbaum, Annu Rev Physiol 2003, 65, 453, 10.1146/annurev.physiol.65.092101.142734; b) R. Gauss, R. Seifert, U. B. Kaupp, Nature 1998, 393 (6685), 583, 10.1038/31248.

[6] a) J. Kusch, V. Nache, K. Benndorf, J Physiol 2004, 560 (Pt 3), 605, 10.1113/jphysiol.2004.070193; b) M. Holmgren, J Gen Physiol 2003, 121 (1), 61, 10.1085/jgp.20028722.

[7] a) K. Benndorf, D. DiFrancesco, Proc Natl Acad Sci U S A 2024, 121 (16), e2400523121, 10.1073/pnas.2400523121; b) K. Benndorf, U. Enke, D. Tewari, J. Kusch, H. Liu, H. Sun, R. Schmauder, C. Sattler, Proc Natl Acad Sci U S A 2025, 122 (5), e2422533122, 10.1073/pnas.2422533122.

[8] a) M. Li, X. Zhou, S. Wang, I. Michailidis, Y. Gong, D. Su, H. Li, X. Li, J. Yang, Nature 2017, 542 (7639), 260, 10.1038/nature20819; b) X. Zheng, Z. Fu, D. Su, Y. Zhang, M. Li, Y. Pan, H. Li, S. Li, R. A. Grassucci, Z. Ren, Z. Hu, X. Li, M. Zhou, G. Li, J. Frank, J. Yang, Nat Struct Mol Biol 2020, 27 (7), 625, 10.1038/s41594-020-0433-5; c) J. Xue, Y. Han, W. Zeng, Y. Jiang, Neuron 2022, 110 (1), 86, 10.1016/j.neuron.2021.10.006; d) Z. Hu, J. Yang, Channels (Austin) 2023, 17 (1), 2273165, 10.1080/19336950.2023.2273165; e) Z. M. James, A. J. Borst, Y. Haitin, B. Frenz, F. DiMaio, W. N. Zagotta, D. Veesler, Proc Natl Acad Sci U S A 2017, 114 (17), 4430, 10.1073/pnas.1700248114; f) Z. Hu, X. Zheng, J. Yang, Nat Commun 2023, 14 (1), 4284, 10.1038/s41467-023-39971-8; g) C. H. Lee, R. MacKinnon, Cell 2019, 179 (7), 1582, 10.1016/j.cell.2019.11.006; h) Cell 2017, 168 (1-2), 111, 10.1016/j.cell.2016.12.023; i) V. Burtscher, J. Mount, J. Huang, J. Cowgill, Y. Chang, K. Bickel, J. Chen, P. Yuan, B. Chanda, Nat Commun 2024, 15 (1), 5216, 10.1038/s41467-024-49599-x; j) A. Saponaro, D. Bauer, M. H. Giese, P. Swuec, A. Porro, F. Gasparri, A. S. Sharifzadeh, A. Chaves-Sanjuan, L. Alberio, G. Parisi, G. Cerutti, O. B. Clarke, K. Hamacher, H. M. Colecraft, F. Mancia, W. A. Hendrickson, S. A. Siegelbaum, D. DiFrancesco, M. Bolognesi, G. Thiel, B. Santoro, A. Moroni, Mol Cell 2021, 81 (14), 2929, 10.1016/j.molcel.2021.05.033; k) A. Saponaro, J. H. Krumbach, A. Chaves-Sanjuan, A. S. Sharifzadeh, A. Porro, R. Castelli, K. Hamacher, M. Bolognesi, D. DiFrancesco, O. B. Clarke, G. Thiel, A. Moroni, Proc Natl Acad Sci U S A 2024, 121 (27), e2402259121, 10.1073/pnas.2402259121; l) B. Yu, Q. Lu, J. Li, X. Cheng, H. Hu, Y. Li, T. Che, Y. Hua, H. Jiang, Y. Zhang, C. Xian, T. Yang, Y. Fu, Y. Chen, W. Nan, P. J. McCormick, B. Xiong, J. Duan, B. Zeng, Y. Li, Y. Fu, J. Zhang, J Biol Chem 2024, 300 (6), 107288, 10.1016/j.jbc.2024.107288.

[9] M. G. Derebe, D. B. Sauer, W. Zeng, A. Alam, N. Shi, Y. Jiang, Proc Natl Acad Sci U S A 2011, 108 (2), 598, 10.1073/pnas.1013636108.

[10] J. Xue, Y. Han, W. Zeng, Y. Wang, Y. Jiang, Neuron 2021, 109 (8), 1302, 10.1016/j.neuron.2021.02.007.

[11] W. Kopec, B. S. Rothberg, B. L. de Groot, Nat Commun 2019, 10 (1), 5366, 10.1038/s41467-019-13227-w.

[12] a) D. Bauer, J. Wissmann, A. Moroni, G. Thiel, K. Hamacher, Function (Oxf) 2022, 3 (3), zqac019, 10.1093/function/zqac019; b) J. H. Krumbach, D. Bauer, A. S. Sharifzadeh, A. Saponaro, R. Lautenschlager, K. Lange, O. Rauh, D. DiFrancesco, A. Moroni, G. Thiel, K. Hamacher, J Gen Physiol 2023, 155 (10), 10.1085/jgp.202313364.

[13] S. Ahrari, T. N. Ozturk, N. D’Avanzo, Biophys J 2022, 121 (11), 2206, 10.1016/j.bpj.2022.04.024.

[14] A. Elbahnsi, J. Cowgill, V. Burtscher, L. Wedemann, L. Zeckey, B. Chanda, L. Delemotte, Elife 2023, 12, 10.7554/eLife.80303.

[15] V. Burtscher, L. Wang, J. Cowgill, Z. W. Chen, C. Edge, E. Smith, Y. Chang, L. Delemotte, A. S. Evers, B. Chanda, Sci Adv 2025, 11 (1), eadr7427, 10.1126/sciadv.adr7427.

[16] H. Liu, J. Biedermann, H. Sun, Commun Biol 2025, 8 (1), 1272, 10.1038/s42003-025-08705-5.

[17] Y. K. Aldakul, M. Schewe, C. Coll-Diez, S. Hwang, T. Baukrowitz, H. Sun, Sci Adv 2025, 11 (32), eadx1680, 10.1126/sciadv.adx1680.

[18] B. Zadek, C. M. Nimigean, J Gen Physiol 2006, 127 (6), 673, 10.1085/jgp.200609534.

[19] O. S. Smart, J. G. Neduvelil, X. Wang, B. A. Wallace, M. S. Sansom, J Mol Graph 1996, 14 (6), 354, 10.1016/s0263-7855(97)00009-x.

[20] T. A. Hilder, B. Corry, S. H. Chung, J Biol Phys 2014, 40 (2), 109, 10.1007/s10867-013-9338-4.

[21] a) C. Shi, Y. He, K. Hendriks, B. L. de Groot, X. Cai, C. Tian, A. Lange, H. Sun, Nat Commun 2018, 9 (1), 717, 10.1038/s41467-018-03179-y; b) R. N. Roy, K. Hendriks, W. Kopec, S. Abdolvand, K. L. Weiss, B. L. de Groot, A. Lange, H. Sun, L. Coates, IUCrJ 2021, 8 (Pt 3), 421, 10.1107/S205225252100213X; c) S. Minniberger, S. Abdolvand, S. Braunbeck, H. Sun, A. J. R. Plested, J Mol Biol 2023, 435 (6), 167970, 10.1016/j.jmb.2023.167970; d) M. J. Gallenito, M. T. Clabbers, J. Lin, J. Hattne, T. Gonen, Adv Sci (Weinh) 2025, 12 (30), e04881, 10.1002/advs.202504881.

[22] W. Kopec, D. A. Kopfer, O. N. Vickery, A. S. Bondarenko, T. L. C. Jansen, B. L. de Groot, U. Zachariae, Nat Chem 2018, 10 (8), 813, 10.1038/s41557-018-0105-9.

[23] S. Ye, Y. Li, Y. Jiang, Nat Struct Mol Biol 2010, 17 (8), 1019, 10.1038/nsmb.1865.

[24] a) A. Acharya, K. Jana, D. Gurvic, U. Zachariae, U. Kleinekathofer, Biophys J 2023, 122 (14), 2996, 10.1016/j.bpj.2023.03.035; b) J. Biedermann, S. Braunbeck, A. J. R. Plested, H. Sun, Proc Natl Acad Sci U S A 2021, 118 (8), 10.1073/pnas.2012843118.

[25] Schrodinger, LLC, unpublished.

[26] J. Huang, S. Rauscher, G. Nawrocki, T. Ran, M. Feig, B. L. de Groot, H. Grubmuller, A. D. MacKerell, Jr., Nat Methods 2017, 14 (1), 71, 10.1038/nmeth.4067.

[27] W. L. Jorgensen, J. Chandrasekhar, J. D. Madura, R. W. Impey, M. L. Klein, Journal of Chemical Physics 1983, 79 (2), 926, 10.1063/1.445869.

[28] S. Jo, T. Kim, V. G. Iyer, W. Im, J Comput Chem 2008, 29 (11), 1859, 10.1002/jcc.20945.

[29] a) B. Roux, Biophys J 2008, 95 (9), 4205, 10.1529/biophysj.108.136499; b) J. Gumbart, F. Khalili-Araghi, M. Sotomayor, B. Roux, Biochim Biophys Acta 2012, 1818 (2), 294, 10.1016/j.bbamem.2011.09.030; c) C. Caleman, D. van der Spoel, Angew Chem Int Ed Engl 2008, 47 (8), 1417, 10.1002/anie.200703987.

[30] C. Kutzner, H. Grubmuller, B. L. de Groot, U. Zachariae, Biophysical Journal 2011, 101 (4), 809, 10.1016/j.bpj.2011.06.010.

[31] D. P. Tieleman, H. J. C. Berendsen, The Journal of Chemical Physics 1996, 105 (11), 4871, 10.1063/1.472323.

[32] M. J. Abraham, T. Murtola, R. Schulz, S. Páll, J. C. Smith, B. Hess, E. Lindahl, SoftwareX 2015, 1-2, 19, 10.1016/j.softx.2015.06.001.

[33] T. Darden, D. York, L. Pedersen, The Journal of Chemical Physics 1993, 98 (12), 10089, 10.1063/1.464397.

[34] G. Bussi, D. Donadio, M. Parrinello, J Chem Phys 2007, 126 (1), 014101, 10.1063/1.2408420.

[35] M. Parrinello, A. Rahman, Journal of Applied Physics 1981, 52 (12), 7182, 10.1063/1.328693.

[36] B. Hess, H. Bekker, H. J. C. Berendsen, J. G. E. M. Fraaije, Journal of Computational Chemistry 1997, 18 (12), 1463, 10.1002/(sici)1096-987x(199709)18:12<1463::Aid-jcc4>3.0.Co;2-h.

[37] N. Michaud-Agrawal, E. J. Denning, T. B. Woolf, O. Beckstein, J Comput Chem 2011, 32 (10), 2319, 10.1002/jcc.21787.

[38] C. R. Harris, K. J. Millman, S. J. van der Walt, R. Gommers, P. Virtanen, D. Cournapeau, E. Wieser, J. Taylor, S. Berg, N. J. Smith, R. Kern, M. Picus, S. Hoyer, M. H. van Kerkwijk, M. Brett, A. Haldane, J. F. Del Rio, M. Wiebe, P. Peterson, P. Gerard-Marchant, K. Sheppard, T. Reddy, W. Weckesser, H. Abbasi, C. Gohlke, T. E. Oliphant, Nature 2020, 585 (7825), 357, 10.1038/s41586-020-2649-2.

[39] J. D. Hunter, Computing in Science & Engineering 2007, 9 (3), 90, 10.1109/mcse.2007.55.

[40] W. McKinney, in 2011, 1–9.

[41] P. Virtanen, R. Gommers, T. E. Oliphant, M. Haberland, T. Reddy, D. Cournapeau, E. Burovski, P. Peterson, W. Weckesser, J. Bright, S. J. van der Walt, M. Brett, J. Wilson, K. J. Millman, N. Mayorov, A. R. J. Nelson, E. Jones, R. Kern, E. Larson, C. J. Carey, I. Polat, Y. Feng, E. W. Moore, J. VanderPlas, D. Laxalde, J. Perktold, R. Cimrman, I. Henriksen, E. A. Quintero, C. R. Harris, A. M. Archibald, A. H. Ribeiro, F. Pedregosa, P. van Mulbregt, C. SciPy, Nat Methods 2020, 17 (3), 261, 10.1038/s41592-019-0686-2.

[42] M. B. Ulmschneider, C. Bagneris, E. C. McCusker, P. G. Decaen, M. Delling, D. E. Clapham, J. P. Ulmschneider, B. A. Wallace, Proc Natl Acad Sci U S A 2013, 110 (16), 6364, 10.1073/pnas.1214667110.

[43] W. Humphrey, A. Dalke, K. Schulten, J Mol Graph 1996, 14 (1), 33, 10.1016/0263-7855(96)00018-5.

[44] K. Benndorf, in (Ed.: B. N. Sakmann, E.), 2nd edition, Springer, 1995.

[45] V. Nache, J. Kusch, V. Hagen, K. Benndorf, Biophys J 2006, 90 (9), 3146, 10.1529/biophysj.105.078667.

