## Supplementary Information for "Selectivity Filter Dynamics Define Ion Conductance and Selectivity Differences in CNG and HCN Channels"

**Table of Content**

Supplementary Figures
Supplementary Tables
Supplementary References

### Supplementary Figures

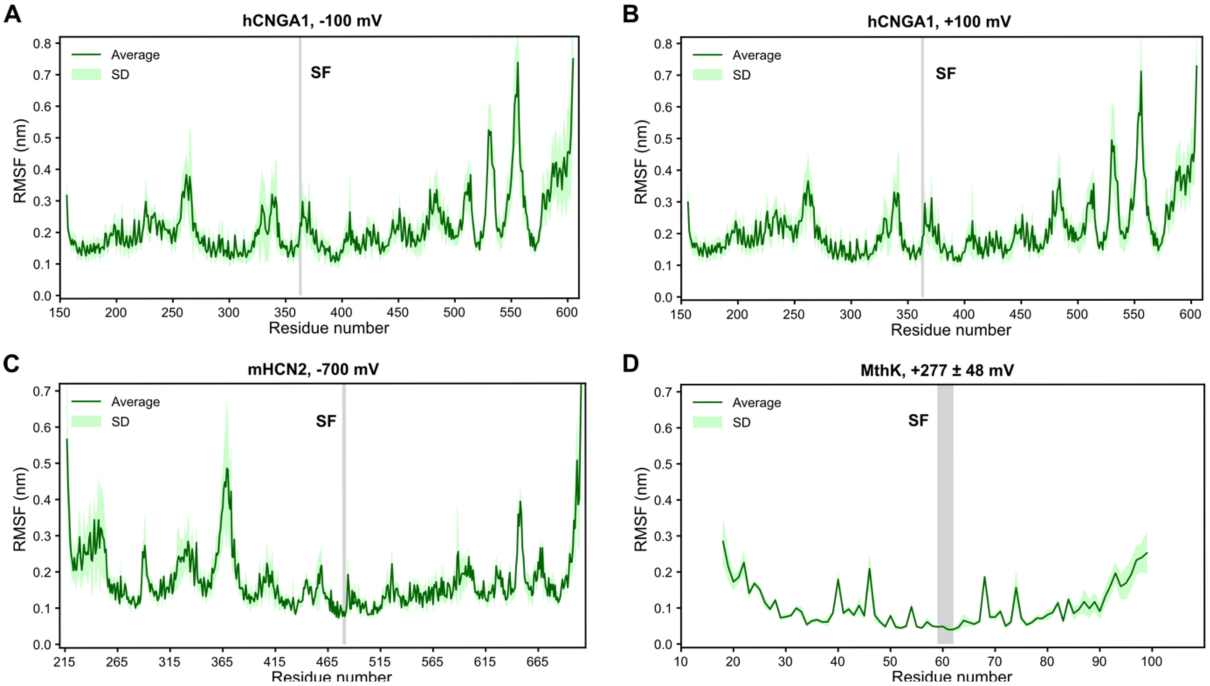


**Supplementary Figure S1 | Root-mean-square-fluctuation (RMSF).** For each MD simulation setup, the green line represents the average RMSF value from five runs, while the light green shading indicates the standard deviation. The grey-highlighted region marks the selectivity filter (SF). **A** Simulations of the full length hCNGA1 channel with $K^+$ under -100 mV transmembrane voltage. **B** Simulations of the full length hCNGA1 channel with $K^+$ under +100 mV transmembrane voltage. **C** Simulations of the full length mHCN2 channel with $K^+$ under -700 mV transmembrane voltage. **D** Simulations of the transmembrane pore domain of MthK channel with $K^+$ under -277±48 mV transmembrane voltage.

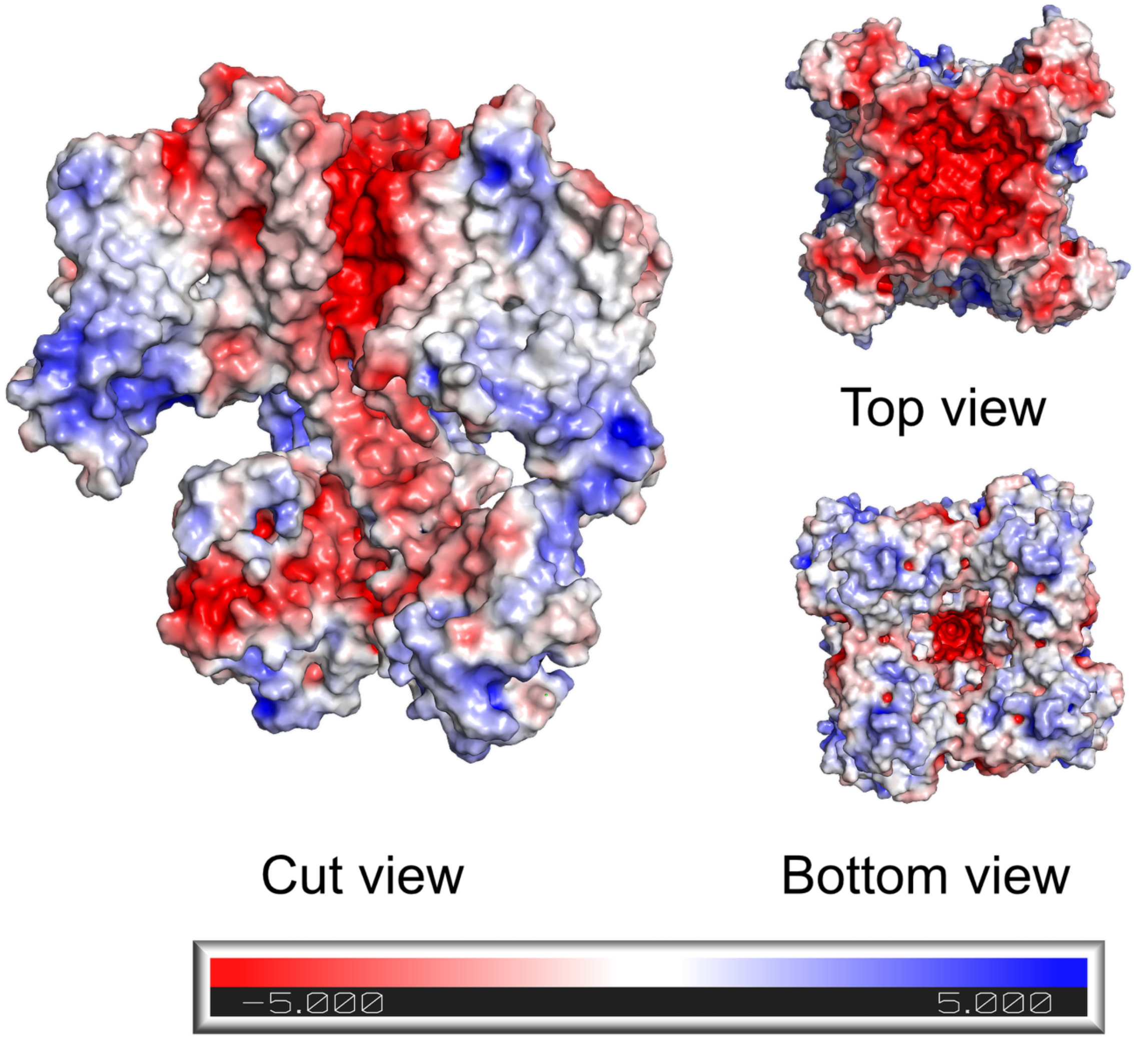


**Supplementary Figure S2 | Electrostatic surface of hCNGA1 channel.** (left) Electrostatic surface representation of the hCNGA1 channel viewed from the interior. (Top right) Top view from extracellular side. (Bottom right) Bottom view from intracellular side. Negatively charged regions are shown in red, and positively charged regions in blue.

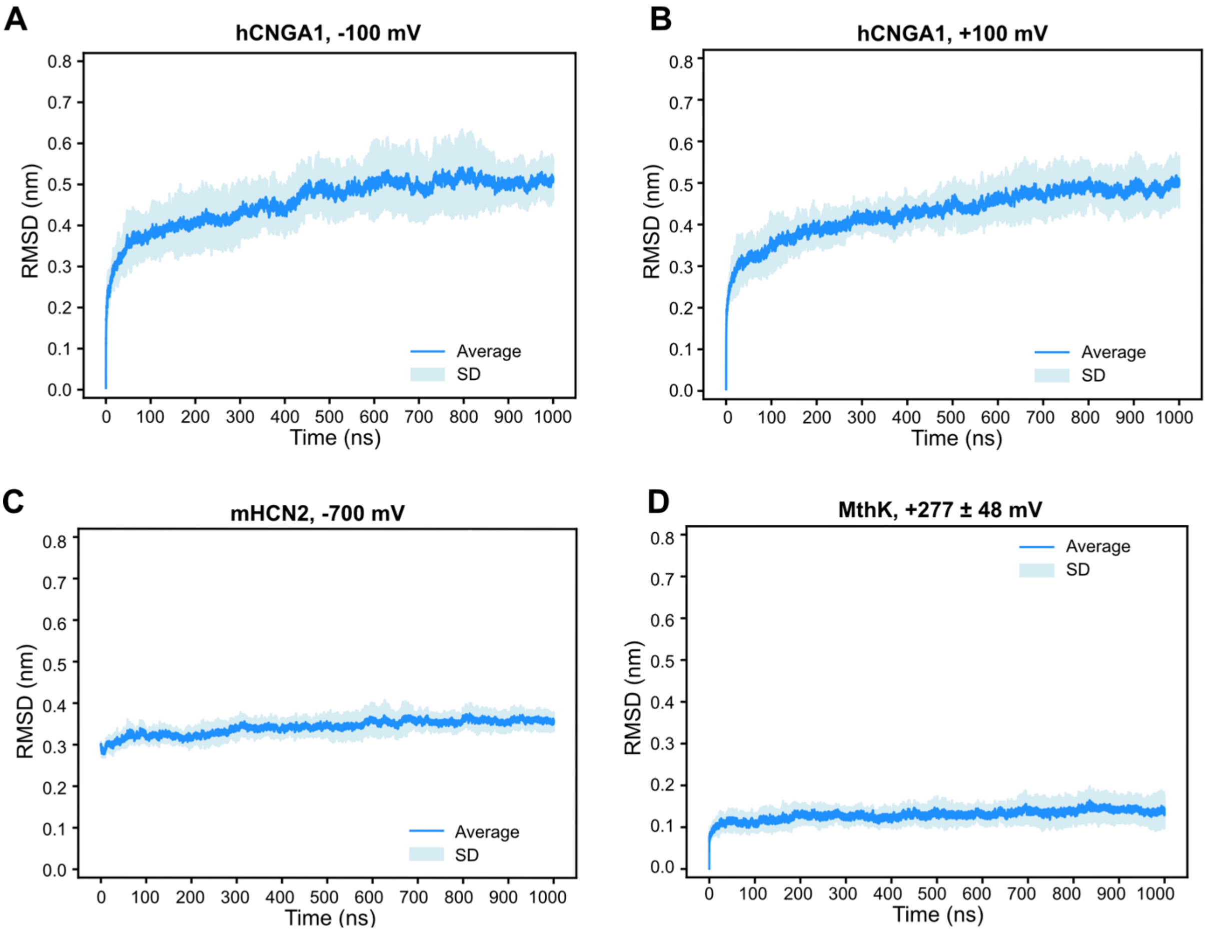


**Supplementary Figure S3 | Root-mean-square-deviation (RMSD).** For each MD simulation setup, the blue line represents the average RMSD value from five runs, with the blue shading indicating the standard deviation. **A** Simulations of the full length hCNGA1 channel with $K^+$ under -100 mV transmembrane voltage. **B** Simulations of the full length hCNGA1 channel with $K^+$ under +100 mV transmembrane voltage. **C** Simulations of the full length mHCN2 channel with $K^+$ under -700 mV transmembrane voltage. **D** Simulations of the transmembrane pore domain of MthK channel with $K^+$ under -277±48 mV transmembrane voltage.

### Supplementary Tables

**Supplementary Table S1 | List of simulated systems.** Simulations of hCNGA1 channel and mHCN2 channel started with full length structures and simulations of MthK channel started with only the transmembrane pore domain. All of the simulations were performed with a temperature at 300 K. * Simulations using computational electrophysiology method[1]; others using an external electric field.

| Simulation system | Force field | Cation | Concentration (mM) | Time (ns) | V (mV) | Replicates | Mean ion permeations | γ (pS) |
|---|---|---|---|---|---|---|---|---|
| hCNGA1 | CHARMM36m | $K^+$ | 150 | 1000 | +100 | 5 | 41 ± 13 | 66 ± 20 |
| hCNGA1 | CHARMM36m | $K^+$ | 150 | 1000 | -100 | 5 | 76 ± 52 | 122 ± 84 |
| mHCN2 | CHARMM36m | $K^+$ | 900 | 1000 | -700 | 5 | 12 ± 3 | 2.7 ± 0.7 |
| MthK * | CHARMM36m | $K^+$ | 600 | 3000 | +277 ± 48 | 5 | 28 ± 16 | 5.3 ± 3.2 |

**Supplementary Table S2 | Simulation system details.** * Simulations using computational electrophysiology method[1]; others using an external electric field.

| Simulation system | Cation | Number of protein atoms | Number of water molecules | Number of POPC | Number of cations | Number of $Cl^-$ | System atoms in total |
|---|---|---|---|---|---|---|---|
| hCNGA1, -100 mV | $K^+$ | 29760 | 42563 | 319 | 109 | 109 | 200413 |
| hCNGA1, +100 mV | $K^+$ | 29760 | 42563 | 319 | 109 | 109 | 200413 |
| mHCN2 | $K^+$ | 32296 | 69920 | 437 | 1079 | 1087 | 302780 |
| MthK * | $K^+$ | 10424 | 12028 | 284 | 146 | 122 | 84832 |

**Supplementary References**

[1] C. Kutzner, H. Grubmuller, B. L. de Groot, U. Zachariae, *Biophysical Journal* **2011**, *101* (4), 809, https://doi.org/10.1016/j.bpj.2011.06.010.
